# Automated Spatially Targeted Optical Micro Proteomics (autoSTOMP) to determine protein complexity of subcellular structures

**DOI:** 10.1101/783340

**Authors:** Bocheng Yin, Roberto Mendez, Xiaoyu Zhao, Rishi Rahkit, Ku-Lung Hsu, Sarah E Ewald

**Affiliations:** Department of Microbiology, Immunology and Cancer Biology and the Carter Immunology Center, University of Virginia School of Medicine Charlottesville VA, USA; Department of Chemistry, University of Virginia, Charlottesville, VA, USA; Mitokinin Inc, 953 Indiana St, San Francisco, CA

## Abstract

Spatially Targeted Optical Micro Proteomics (STOMP) is a method to study region-specific protein complexity of a biological specimen. STOMP uses a confocal microscope to both visualize structures of interest and to tag the proteins within those structures by a photo-driven crosslinking reaction so that they can be affinity purified and identified by mass spectrometry^1^. STOMP has the potential to perform discovery proteomics on sub-cellular structures in a wide range of primary cells types and biopsy-scale tissue samples. However, two significant limitations have prevented the broad adoption of this technique by the scientific community. First, STOMP is performed across two software platforms written in different languages, which requires user operation at each field of view. Up to 48 hours of microscope time is necessary to tag sufficient protein (∼1 μg) for mass spectrometry making STOMP prohibitively time and labor-consuming for many researchers. Second, the original STOMP protocol uses a custom photo-crosslinker that limits the accessibility of the technique for some user. To liberate the user, we developed a protocol that automates communication between Zeiss Zen Black imaging software and FIJI image processing software using a customizable code in SikuliX. To fully automate STOMP (autoSTOMP), this protocol includes a tool to make tile array, autofocus and capture images of fields of view across the sample; as well as a method to modify the file that guides photo-tagging so that subsets of the structures of interest can be targeted. To make this protocol broadly accessible, we implemented a commercially available biotin-benzophenone crosslinker as well as a procedure to block endogenous biotin and purify tagged proteins using magnetic streptavidin beads. Here we demonstrate that autoSTOMP can efficiently label, purify and identify proteins that belong to structures measuring 1-2 μm in diameter using human foreskin fibroblasts or mouse bone marrow-derived dendritic cells infected with the protozoan parasite *Toxoplasma gondii* (*Tg*). The autoSTOMP platform can easily be adapted to address a range of research questions using Zeiss Zen Black microscopy systems and LC-MS protocols that are standard in many institutional research cores.

## INTRODUCTION

Localization is a critical regulatory component of protein function. Currently, there is a tremendous demand for biochemical and microscopy techniques to assess regional protein complexity and function in tissues, cells, and organelles. Spatially targeted optical micro proteomics (STOMP) was designed to identify the proteome of regional structures.^1^ STOMP was first implemented to identify components of amyloid plaques in brain sections from a mouse model of prion disease and Alzheimer’s disease patients.^1^ In STOMP, structures of interest (SOI) are stained for fluorescence microscopy and identified by confocal imaging. Confocal images are used to generate a “MAP” file. Using a custom macro, the “MAP” file guides the 2-photon laser to revisit structures of interest and conjugate bifunctional, UV-activated affinity purification tags to proteins within those structures. The tagged proteins are affinity-purified for identification by liquid chromatography-mass spectrometry (LC-MS). STOMP can be performed on a wide range of SOI where one or more labeling reagents (e.g., antibodies, stains) are available to visualize it^2^. This makes STOMP well suited to perform discovery proteomics on sub-cellular structures in a wide range of primary cells types and biopsy-scale tissue samples.

Proximity biotinylation methods like BioID and APEX, require a biotin-targeting protein that is genetically engineered and stably expressed in a cell, so these tools are largely limited to cell lines or cells that can be easily transduced.^2^ In comparison, STOMP can be performed on a wider range of structures of primary cells and tissues. In proximity biotinylation, regional selectivity can be mediated by expressing the biotinylation enzyme on a specific side of a membrane-spanning protein (e.g., PPOX, the mitochondrial matrix protein), by changing the length of the enzyme-linker domain or by changing the duration of the biotinylation reaction.^3^ However, if there is more than one pool of the label-targeting protein in the cell (e.g., ER and plasma membrane), all neighboring proteins will be labeled. In comparison, STOMP uses a microscope image to guide tagging, so the user has the potential to customize the photo-crosslinking coordinates to select only one pool for labeling. For example, by dilating, eroding, or finding the edges of an SOI, protein belonging to the core structure or near-neighbor structures can be identified. Alternatively, co-localization stains can be used to identify and tag the subpopulations of the SOI. In this way, region-specific SOI proteomes can be determined to inform hypotheses about the biology regulated by candidate proteins. In antibody-mediated proximity biotinylation (e.g., SPPLAT or BAR) a labeling enzyme is attached to an antibody specific to a protein of interest; this facilitates investigating a wide range of biological specimens at a 10-100 nm resolution similar to APEX or BioID.^2,4^ However, its regional selectivity is limited compared to STOMP. Laser capture microdissection is often used to isolate intact cells for omics analysis.^5^ Like STOMP laser capture microdissection provides a high degree of spatial selectivity. However, it is a tedious process with limited resolution beyond 10 μm compared to STOMP which approaches 1 μm resolution.^1,5^

The original STOMP protocol took advantage of a bi-functional 6-HIS-benzophenone crosslinker for UV-mediated SOI protein tagging. The benzophenone moiety attacks any local carbon or nitrogen upon exposure to UV-wavelength light. The HIS tag was implemented because it could be synthesized in a cost-efficient way and benzophenone-HIS proteins were easily eluted from nickel agarose columns. However, a chemical treatment of the sample with diethyl pyrocarbonate (DEPC) was necessary to reduce background binding of endogenous histidine to the nickel agarose matrix. A limitation of this strategy was that atypical histidine modification by DEPC could negatively impact the identification of some peptides by LC-MS analysis^67^. Moreover, researchers that did not have access to peptide synthesis tools may not have been able to easily access the 6-HIS-benzophenone reagent to perform STOMP.

A second major shortcoming of the original STOMP protocol was that the user must manually process images in Zeiss Zen black and FIJI at each field of view. Approximately 48 hours of laser time were required to tag sufficient protein (1-2 μg) for MS analysis. Depending on the size of the SOI, this required a user to operate the programs every 20-90 minutes for the duration of the photo-labeling process^1^. The benefits of this approach were that it provided tremendous flexibility in selecting biotinylation targets, and it allowed the user to assess each image for staining artifacts. However, extensive user intervention also introduced the opportunity for user error or experimental variability.

Our goal was to develop a fully automated protocol for STOMP using commercially available reagents and an intuitive user interface that could be implemented using hardware standard in many labs and research cores. We refer to this protocol as automated STOMP (autoSTOMP). To communicate between Zeiss Zen Black and FIJI software, we used the open-source, icon recognition software SikuliX (http://sikulix.com). SikuliX is written in the Jython (https://www.jython.org) language and can be easily coded in by users without extensive programming skills. The autoSTOMP protocol integrates an autofocus and tile array platform that eliminates the need for time-consuming user intervention. Finally, we implemented a new biochemistry protocol using biotin-benzophenone that includes steps to block endogenous biotin and streptavidin-precipitation enrichment of biotinylated proteins. To validate the specificity of autoSTMOP laser targeting and biotinylated protein enrichment we infected human or murine cells with the protozoan parasite *Toxoplasma gondii (Tg)*.^8–10^ *Tg* has a diverse proteome of over 6,000 predicted protein-coding genes that are differentially expressed across three parasite life stages. Protozoa diverged from the common mammalian ancestor approximately 2 billion years ago, so the *Tg* proteome is sufficiently distinct from mouse and human proteomes to facilitate peptide alignment for the majority of proteins. These proof of principle experiments establish autoSTOMP as a robust and reproducible protocol to identify regional proteomes in biological samples.

## MATERIALS AND METHODS

### Reagents and consumables

Unless otherwise noted chemical reagents and consumables were purchased from Thermo Fisher Scientific, USA, and used according to manufacturer instructions.

### *Tg* infections and immunofluorescence staining

*Tg* is cultured and stored under BSL2 conditions in accordance with the University of Virginia Environmental Health and Safety approved Biosafety Protocol. The Type II *Tg* parasite strain ME49 was used in all experiments. *Tg* was passaged in confluent human foreskin fibroblasts (HFF) grown in 25cm^2^ tissue culture flasks. Parasites and HFFs were cultured in complete DMEM media containing DMEM (11965118), 10% heat-shocked FBS (12303C-500ML, Sigma Aldrich), 1% Penicillin/Streptomycin solution (15140163, Fisher Scientific), 1% L-glutamine solution (25030164, Fisher Scientific), and 1% 100 mM sodium pyruvate solution (11360070, Life Technologies), 1% 1 M HEPES solution (15630080, Life Technologies). Media was stored at 4 °C and warmed to 37 °C before use. C57BL/6 mice were bred in the University of Virginia vivarium in accordance with ABSL-1 standards and AALAC approved protocol. Mouse bone marrow-derived dendritic cells (mBMDCs) were differentiated for 6-8 days in complete RPMI media supplemented with 10% mouse GMCSF supernatant derived from B16 cells stably expressing mouse GMCSF as previously described^8^. THP-1 cells were cultured in suspension in complete RPMI media containing RPMI (10-040-CV, Corning), 10% heat-shocked FBS, 1% Penicillin/Streptomycin solution, 1% L-glutamine solution, and 1% 100 mM sodium pyruvate solution, 1% 1 M HEPES.

For experimental infections, 0.5×10^6^ THP-1 cells were differentiated with complete RPMI media containing 100 ng/mL PMA (AAJ63916MCR, Fisher Scientific) for 2 days on 12 mm cover glass (64-0712, Harvard Apparatus, USA) coated with poly-D-lysine (ICN15017550). mBMDCs were plated 1.35×10^6^ per well of a 12 well plate containing a single 18 mm round cover glass (64-0714, Harvard Apparatus, USA) coated with poly-D-lysine. HFFs were grown to confluency on 18mm round cover glass. *Tg* was grown in HFFs until large intracellular parasite vacuoles were observed. Intracellular *Tg* was harvested by scraping infected HFF monolayers with a rubber policeman and syringe lysing the HFFs through a 25G blunt-ended syringe. *Tg* was counted on a hemocytometer and added to host cells at a multiplicity of infection (MOI) of 10, or for HFF infections, 8.8×10^4^ parasites per mm^2^. The *Tg* infected HFFs on coverslips were harvested after 2 hours.

### Slide Preparation for STOMP

Coverslips were fixed with 100% methanol at −20 to −30 °C (prechilled) for 15 min. Methanol was decanted and samples were washed three times with room temperature PBS. If staining did not take place immediately, PBS was aspirated and coverslips were stored at −30 °C. To stain *Tg*, the slides were blocked with 2% BSA in TBS at room temperature for 1 hour, a rabbit polyclonal antibody specific to soluble tachyzoite antigen (STAG) directly conjugated to FITC (PA1-7253, Invitrogen) was diluted 1:300 in TBS-T (0.1% Tween20) and coverslips were stained at room temperature for 1 hour. Slides were washed three times with TBST. Endogenous biotin was blocked with Avidin/Biotin Blocking Kit (SP-2001, Vector Laboratories) following the manufacturer’s protocol. The stock solution of 0.5 M biotin-dPEG®3-benzophenone (biotin-BP, 10267, Quanta BioDesign) in anhydrous DMSO (89139-666, VWR) is stable for 12 months when stored with desiccant in the dark at −30 °C. The biotin-BP mounting media must be prepared fresh (used within 4 hours) by diluting the biotin-BP stock solution in 50/50 (v/v) DMSO/water to a working concentration of 1 mM biotin-BP. Coverslips were mounted in 12 μL biotin-BP mounting media in a cold room (4 °C) with weak light and sealed with nail polish (Insta-Dri Fast Dry Nail and Double Duty Base and Topcoat, Sally Hansen). If multiple slides were necessary, coverslips were mounted the day of crosslinking and stored in the dark at room temperature before microscopy.

### Validation of biotin-benzophenone cross-linking by microscopy

Following biotin-BP cross-linking, each coverslip was soaked in DI water (RT, dark) for 30 minutes and the nail polish seal was gently pulled away. Excess mounting media was removed by three washes with 50/50 (v/v) DMSO/water followed by three washes with Mili-Q water. Slides were incubated in TBST (0.1% Tween20) containing 1: 500 dilution Alexa Fluor® 594 Streptavidin (#016-580-084, Jackson ImmunoResearch Lab) for 45 min. Coverslips were washed three times in TBST (0.1% Tween20) and mounted with mounting media containing DAPI (H-1000, Vector Laboratories) for imaging.

### Image acquisition, mask generation and UV-biotinylation using autoSTOMP

All microscopy and cross-linking were performed on an LSM880 confocal microscope (Carl Zeiss, Inc., Germany) and a Chameleon multiphoton light source (Coherent Inc., USA) in the Ewald Lab at the University of Virginia. Images were acquired using Zen Black (Carl Zeiss). Image modification and MAP file generation were performed in FIJI (FIJI Is Just ImageJ)^11^. Structures of Interest (SOI, *Tg* here) were visualized using the 25x oil emersion lens (LD LCI Plan-Apochromat 25x/0.81 mm Korr DIC M27) with immersion oil (518 F for 30°C, refractive index = 1.518, 444970-9000-000, Carl Zeiss) and the argon laser source (488 nm) with 500-530 nm bandwidth. A 512 × 512 pixel^2^ image was acquired for each field of view. Sikulix version 1.1.4 (http://sikulix.com/) was used to automate tasks between software platforms. Python (version 3.6, www.python.org) and Spyder (version 3.2.8, www.spyder-ide.org) were used to generate a tile array across the slide surface (∼ 500 tiles). Initial images acquired by the autofocus function were compared to manually acquired images to validate the signal to noise ratio. Using the SikuliX automation platform, each field of view was processed as follows. First, the SOI in each field of view were imaged. The .czi file was exported into FIJI where a binary image was created, thresholded and/or “erode”, “dilate” or “find edges” functions were used to create a “MAP file” of the regions to be photo cross-linked. The MAP file is converted to a .txt file which is imported by the STOMP macro in Zen Black. In Zen Black, the STOMP macro directs the Chameleon to deliver 720nm light to each pixel defined in the MAP file. Cross-linking four to five million pixels typically labeled sufficient protein (approximately 1 μg) for mass spectrometry analysis. A step-by-step tutorial of the autoSTOMP protocol and all the source codes are deposited at GitHub (GitHub Inc.). https://github.com/boris2008/Sikulix-automates-a-workflow-performed-in-multiple-software-platforms-in-Windows.git

### Streptavidin precipitation

Once the UV-cross linking is complete for each slide, the coverslip was soaked in DI water (RT, dark) for 30 min and the nail polish seal was gently pulled away. Excess mounting media was eliminated by three washes with 50/50 (v/v) DMSO/water followed by three washes with Mili-Q water. Excess water was aspirated and the coverslip was stored at −30 °C while photo-crosslinking was performed on additional slides.

Our purification protocol for Mass Spectrometry (MS) analysis is modified from *Hadley et al.* 2015^1^. Samples are dissociated from the coverslip in 8 M urea lysis buffer containing 100 mM NaCl, 25 mM Tris, 2% SDS, 0.1% tween 20, 2 mM EDTA, 0.2 mM PMSF, and 1x Roche cOmplete Protease inhibitor tablets (11873580001), benzonase (E1014-25KU), RNase A from bovine pancrease (10109142001). Coverslips were placed on a parafilm (Bemis) membrane with cells facing up and incubated with 50 μL urea lysis buffer at room temperature for 30 min. The coverslip was then rinsed with 100 μL Mili-Q water and followed by another incubation with 50 μL urea lysis buffer at room temperature for 30 min. All of the solutions after lysis were collected and combined as lysate in low protein binding 1.5 mL microcentrifuge tubes. To reduce the nucleic acid-protein complex formation and associated lysate viscosity, benzonase (0.1 μL per 5 × 10^6^ cells) was added and incubated at 37 °C for 30 min followed by RNase (0.5 μL per 5 × 10^6^ cells) treatment at 65 °C for 15 min. The lysate was cooled to room temperature and ready for affinity purification.

Four different buffers were used for the affinity purification: TU (50 mM Tris-Cl, 2 M urea, and 150 mM NaCl at pH=7.4), TUST (50 mM Tris-Cl, 2 M urea, 150 mM NaCl, 0.1% SDS, and 0.1% Tween 20 at pH=7.4), TUB (0.5 mM biotin in TU buffer) and 100 mM NH_4_HCO_3_ buffer. The volume of lysate of each sample was filled up to 1 mL with TUST. 10 μL Pierce™ Streptavidin (SA) Magnetic Beads (88817) 50% slurry was washing with 1 mL TUST once and then added to the lysate and mixed by vortexing. The mixture was incubated on a rotator at room temperature for 1 hour to bind the biotinylated proteins (longer incubation time is not recommended as it will increase unspecific binding). The magnet (DynaMag™-2, Invitrogen) was used to pellet the SA beads. To remove unspecific bound proteins, samples were washed three times with 1 mL TUST, five times with 1 mL TU, 1 time with 1 mL TUB, and three times with 1 mL 100 mM NH_4_HCO_3_ buffer. After each wash, SA beads were pelleted on the magnet for 3 min and the buffer was removed by pipetting while on the magnet. For each wash, the SA beads were mixed by vortexing and incubated for 5 minutes before the SA beads were applied to the magnet again. After all washing, the SA beads were resuspended in 100 μL 100 mM NH_4_HCO_3_ buffer.

### Western blot

To validate *Tg* protein enrichment by western blot, total protein content for each sample was normalized using Pierce™Micro BCA kit (#23235, Pierce). Proteins were separated by SDS-PAGE using Mini Gel Tank tool kit (25977, ThermoFisher) and were transferred to PVDF membrane (1704274, Bio-Rad) using Trans-Blot Turbo Transfer System (Bio-Rad). The PVDF membrane was blocked in 2% milk TBST (0.1% Tween 20) solution for 1 hour at room temperature followed by sequential incubation with antibody diluted in TBST (0.1% Tween 20) and alternating washing with TBST (0.1% Tween 20). Streptavidin-HRP (016-030-084, Jackson ImmunoResearch) was used to probe biotin. Proteins were visualized on ChemiDoc™Touch imaging system (Bio-Rad) with HRP substrate (WBLUF0100, Millipore).

### Sample preparation for LC-MS

After affinity purification, on-bead reduction and alkylation of SA beads were performed in 1 mL 25 mM NH_4_HCO_3_ by adding 25 μL 10 mM DTT at room temperature for 30 min and 25 μL 15 mM iodoacetamide in dark at room temperature for 1h in sequence. To quench excess iodoacetamide, 25 μL 5 mM DDT solution was added and incubated for 10 min. Trypsin/Lys-C (V5072 Promega) was directly spiked into the protein lysate at a ratio of Trypsin/Protein = 1:25. Digestion was continued overnight (∼12 h) at 37 °C. Formic acid was added to a final concentration of 1% (v/v) to stop the trypsinization. SA beads were removed by the magnet and the supernatant was saved. 7.5 nmol Angiotensin/Vasoactive Standard (A9650-1MG/V0131-.1MG, Sigma) was spiked into the peptide digest. The peptide digest was desalted using Pierce™ C18 Tips with binding, washing (with 0.1% formic acid in water), and eluting (with 60% acetonitrile in water) steps by following the manufacturer’s instructions. The sample was dried completely on the SpeedVac concentrator (Model: SPD131DDA-115). The peptide digest was dissolved in 21 μL of LC-MS grade water containing 0.1% formic acid (F0507-100ML, Sigma). The digest was filtered with a 0.65 µm pore-size micro-centrifuge filter before it was loaded to an HPLC auto-injection sample.

### Peptide mass quantification by LC-MS

For the initial validation of autoSTOMP, LC-MS was performed on an linear ion trap mass spectrometer (Thermo Scientific LTQ, Thermo Fisher, USA) coupled with LC (LC-20AD, Shimadzu, USA) in the Hsu lab at the University of Virginia. 1 μL (5%) of the peptide digest was injected to a C18 capillary column (20 cm of 5 μm C18 (YMC*GEL) packed in 360 μm o.d. ×75 μm i.d. fused silica) and desalted with 0.05% acetic acid at a 185 nL/min for 30 min at room temperature. After desalting, a gradient of acetonitrile/0.1 M formic acid was applied for 200 min: solvent A (water + 0.05% acetic acid) and solvent B (80% acetonitrile + 0.05% acetic acid) were applied in the order of 0% B for 5 min, 0-25% B for 111 min, 25-45% B for 35 min, 45-95% B for 1 min, 95% B for 7 min, 95-0% B for 6 min, 0% B for 34 min, and re-equilibration with 0% B for 10 min. The separation column ran at room temperature. The eluted peptides were electro-sprayed into the LTQ MS, which was operated in the positive ion mode with a “top 3” data-dependent acquisition method that consisted of one MS scan (m/z: 350-1800) followed by three MS/MS scans of the most abundant ions recorded in the preceding MS scan. For mass assignment IP2 (the Integrated Proteomics Applications, Inc., USA) using the ProLuCID algorithm was used to search against human protein database (https://www.uniprot.org) and *Tg* protein database (released 41, https://toxodb.org/toxo/) with reversed sequence decoys. The peptide and proteins were identified with false discovery rate (FDR) smaller than 1%, separately. Differences in protein and peptide abundances between the samples were based on MS/MS spectral counting using the COMPARE function in the IP2-Integrated Proteomics Pipeline. The resulting MS2 spectra matches were assembled into protein identifications.

To compare protein enrichment in *Tg* with protein enrichment at the parasite vacuole membrane using the “donut” MAP file function, a Thermo Electron Q Exactive HF-X mass spectrometer system with an Easy Spray ion source connected to a Thermo 75 µm × 15 cm C18 Easy Spray column was used at the University of Virginia Biomolecular Analysis Facility Core. 7 µL (33.3%) of the extract was injected and the peptides eluted from the column by an gradient at a flow rate of 0.3 µL/min at 40 °C for more than 1 hour. Solvent A (water + 0.1% formic acid) and solvent B (80% acetonitrile + 0.1% formic acid) were applied in the order of 2% B for 1.5 min, 2-23% B for 51 min, 23-35% B for 10 min, 35-95% B for 1 min, and 95% B for 5 min. The nanospray ion source was operated at 1.9 kV. The digest was analyzed using a Top10 method with the MS scan set to 120K resolution and HCD scans set to 30K resolution. This mode of analysis produces approximately 25000 MS/MS spectra of ions ranging in abundance over several orders of magnitude. The data were analyzed by database searching using the Sequest search algorithm deployed in Proteome Discoverer™ (Thermo Fisher Scientific Inc. USA) against Uniprot mouse and *Tg* database from toxodb with reversed sequence decoys separately. Protein (FDR < 2%) and peptide identification (FDR< 0.2%) were organized and summarized by Scaffold (Proteome Software, Inc).

### Data Analysis

Common contaminants listed on the MaxQuant website http://www.coxdocs.org/doku.php?id=maxquant:start_downloads.htm have been identified and excluded from our protein data. The proteins possess high similarly (> 90% proteins identification and >80% peptide sequence coverage) between human and *Tg* protein were excluded from downstream analysis using a custom python script “myBlast.py” running on the Rivanna computing server (https://arcs.virginia.edu/rivanna) at the University of Virginia. A Student’s t-test was used to perform pair-wise comparison across the three replicates of each sample with R scripts in Rstudio (version 1.1.456, www.rstudio.com/) with R packages (for examples, readr, xlsx, dplyr, tidyverse) and p-values for each protein was reported. Gene Ontology (GO) and KEGG pathway annotation were added using David Bioinformatics Resources 6.8^12^. The enriched proteins and the statistical summarization were made in R scripts with the table organization and statistical tool packages mentioned above. These custom scripts are deposited at GitHub (GitHub Inc.) https://github.com/boris2008/codes-for-validating-STOMP-with-MS.git. Plots were created with R package “ggplot2 and “ggrepel” in R or GraphPad Prism (version 8.2.1).

## RESULTS AND DISCUSSION

### Automated STOMP Workflow

To establish autoSTOMP, we followed the basic procedure to stain structures of interest (SOI) (Figure 1A), to capture immunofluorescence images in Zeiss Zen Black (Figure 1B); to import images to FIJI and generate “MAP” binary text files (Figure 1C); and to cross-link SOI proteins with 720nm light generated by the two-photon laser source using the STOMP Macro developed by Hadley et al. 2014^1^ (Figure 1D). In this protocol, we used a Zeiss LSM 880 confocal microscope. The original STOMP protocol was implemented to identify novel components of beta-amyloid plaques in mouse and human samples using a Zeiss LSM510 microscope and a 10x water immersion objective with a photo-crosslinking resolution of 0.67×0.67×1.48 μm in x, y, z axis respectively (Figure 1A-E).^1^ To perform STOMP we used a 25x/0.81 mm LD LCI Plan-Apochromat oil emersion lens with a theoretical resolution of 0.38×0.38×2.50 μm in the x, y, z axis (based on spatial resolution equations)^13^. Several major modifications to the initial STOP protocol have been implemented for autoSTOMP (Figure 1, light orange boxes). These include using a commercially available Biotin-dPEG®3-benzophenone (biotin-BP) probe (in place of 6-HIS-BP) and a step to block endogenous biotins after SOI staining (Figure 1F). After crosslinking is complete, samples are dissolved in a urea lysis solution and streptavidin-coated magnetic beads are used to enrich biotinylated proteins in place of the previously published nickel agarose column (Figure 1E). On bead trypsin/LysC protease digestion is used to isolate precipitated peptides for LC-MS. In addition, a tile array function was built using Python codes to identify each field of view across sample, automate focus on SOI and acquire SOI images (Figure 1G). The entire protocol is fully automated using the SikuliX icon recognition software (Figure 1H) to transition between the fields of view (Figure 1G), Zen Black and FIJI (Figure 1B-D) without any user intervention. Progress reports or errors in the automation process are automatically sent to a user Gmail account so that the system can be monitored remotely (Figure 1I).

**Figure 1.**
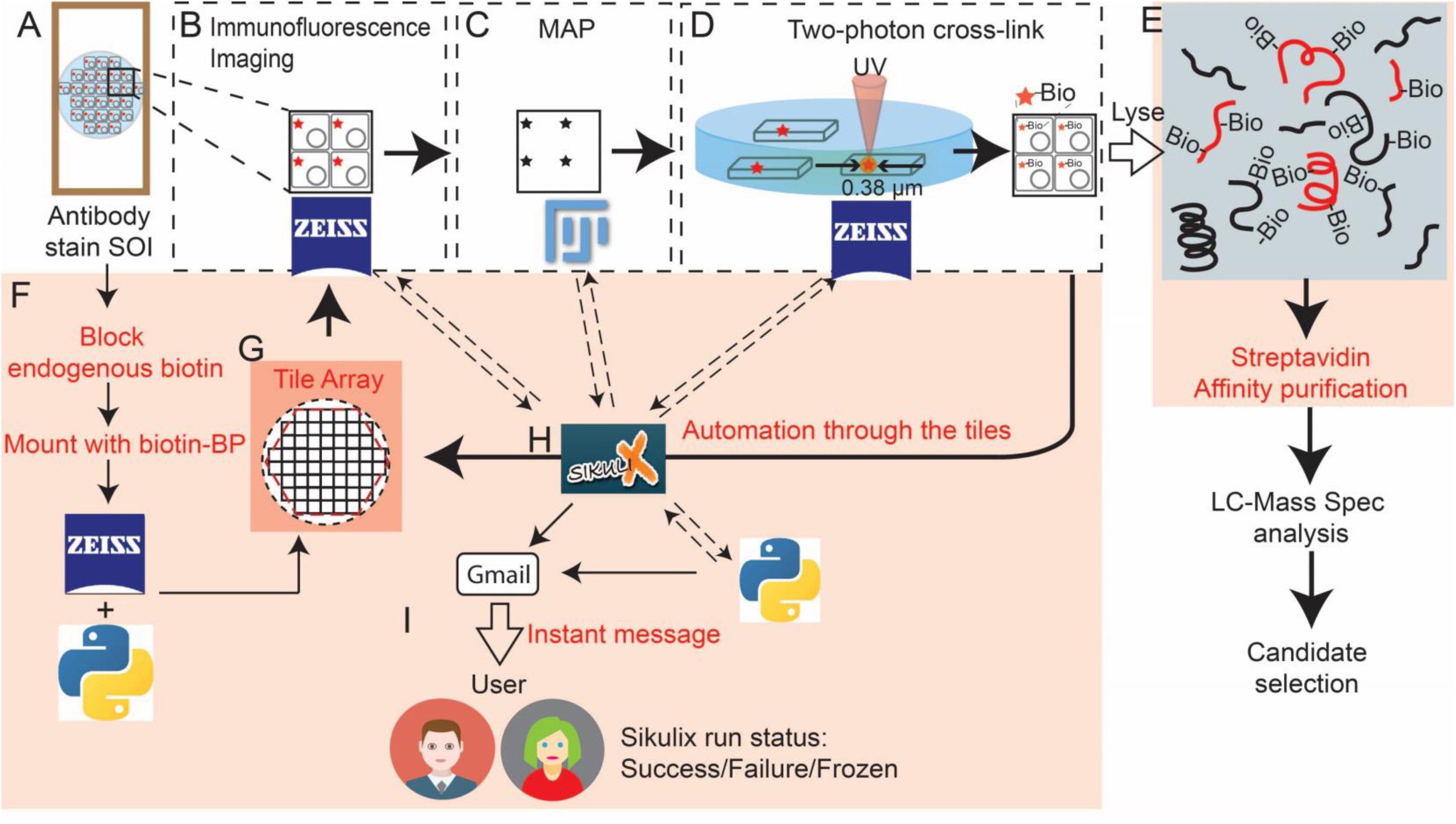
Overview of the Spatially Targeted Optical Micro Proteomics (STOMP) protocol (A-D) and automated-STOMP modification (E-I, orange boxes). **A**, Structures of interest (SOI) are stained for identification by standard fluorescence microscopy. **B**, SOI are identified by immunofluorescence imaging in visible (non-UV) wavelengths using Zeiss Zen Black software. **C**, Immunofluorescence images are exported to FIJI software used to generate a “map” file. **D**, The map file is imported into Zeiss Zen Black using a custom STOMP Macro. The STOMP Macro guides the 2-photon laser to selectively deliver UV energy to each SOI identified in the map, conjugating biotin-BP to any protein within the bounds of each SOI. **E**, Biotinylated SOI proteins are affinity-purified using streptavidin-coated beads and identified by liquid chromatography-mass spectrometry (LC-MS). Both stained (red) and unstained (black) proteins within the SOI are biotinylated. **F**, Slides are mounted in a media containing a bi-functional, biotin-benzophenone (biotin-BP) affinity purification tag that is activated by UV to covalently attach to local carbon or nitrogen. **G**, To automate STOMP (auto-STOMP) a tile array of each field of view across the sample is generated using a custom python script and Zeiss Zen Black coordinate functions. **H**, The SikuliX icon recognition software is used to automate the basic STOMP protocol (auto-STOMP) by integrating the directed network of tasks between Zen Black (autofocus and image acquisition (B) and map guided photo-labeling(C)) and Fiji (image processing and map generation(D)). **I**, Instant messages are delivered to the user through Gmail portal for the update of automation status (Success/Failure/Frozen). Red boxes indicate auto-STOMP protocol updates.

The photo-excitation conditions for autoSTOMP were determined experimentally on a stained macrophage monolayer by modifying the laser power (P), pixel dwell time (D), and the number of iterations (I). Each field of view was targeted then samples were assessed for photobleaching and sample overheating (Figure S-1A). Several conditions cause noticeable photo-bleaching but no overheating (Figure S-1A, red box). P/I/D =3/1/1 was chosen to minimize the total time necessary to photo-crosslink the sample (with optimal photobleaching and without overheating, Figure S-1B). To identify the minimum concentration of biotin-BP necessary to achieve robust SA-594 signal, the concentration of biotin-BP in the mounting media was titrated. Thus, optimal photo-active biotinylation was selected (Figure S-2).

### AutoSTOMP selectively biotinylates SOI

To determine if autoSTOMP can effectively enrich proteins from sub-cellular SOI we infected human foreskin fibroblasts (HFFs) with the ME49 strain of Type II *Toxoplasma gondii (Tg)* which forms intracellular vacuole measuring 1-6 μm (Figure 2A). After two hours of infection, the coverslips were fixed and stained with an antibody raised against soluble *Tg* antigen (commonly referred to as STAG) directly conjugated to FITC (α-*Tg*-488) to identify *Tg* as the SOIs (Figure 2B). A MAP file was generated and used to guide UV-excitation and biotinylation reaction using the autoSTOMP platform (Figure 1). Following UV-crosslinking, coverslips were removed and stained with streptavidin directly conjugated to Alexafluor594 (SA-594) and reimaged (Figure 2D). Co-localization was confirmed by re-imaging the *Tg-*specific antibody in the 488 nm channel (Figure 2E-F). Co-localization of the α-*Tg* signal and SA-594 signal confirmed the accuracy of UV targeting and the efficiency of biotinylation. As expected, the α-*Tg*-488 signal was slightly photobleached upon re-imaging compared to that is before UV-targeting by autoSTOMP (Figure 2E versus 2B). Tile imaging around the field of view identified in the MAP file (Figure 2G, dotted white box) confirmed that autoSTOMP selectively biotinylated SOI identified in the MAP and did not increase background SA-594 signal relative to not-targeted fields of view (Figure 2H). Based on these data, we conclude that autoSTOMP efficiently targets biotin-BP conjugation to regions defined as SOI by immunofluorescence imaging.

**Figure 2.**
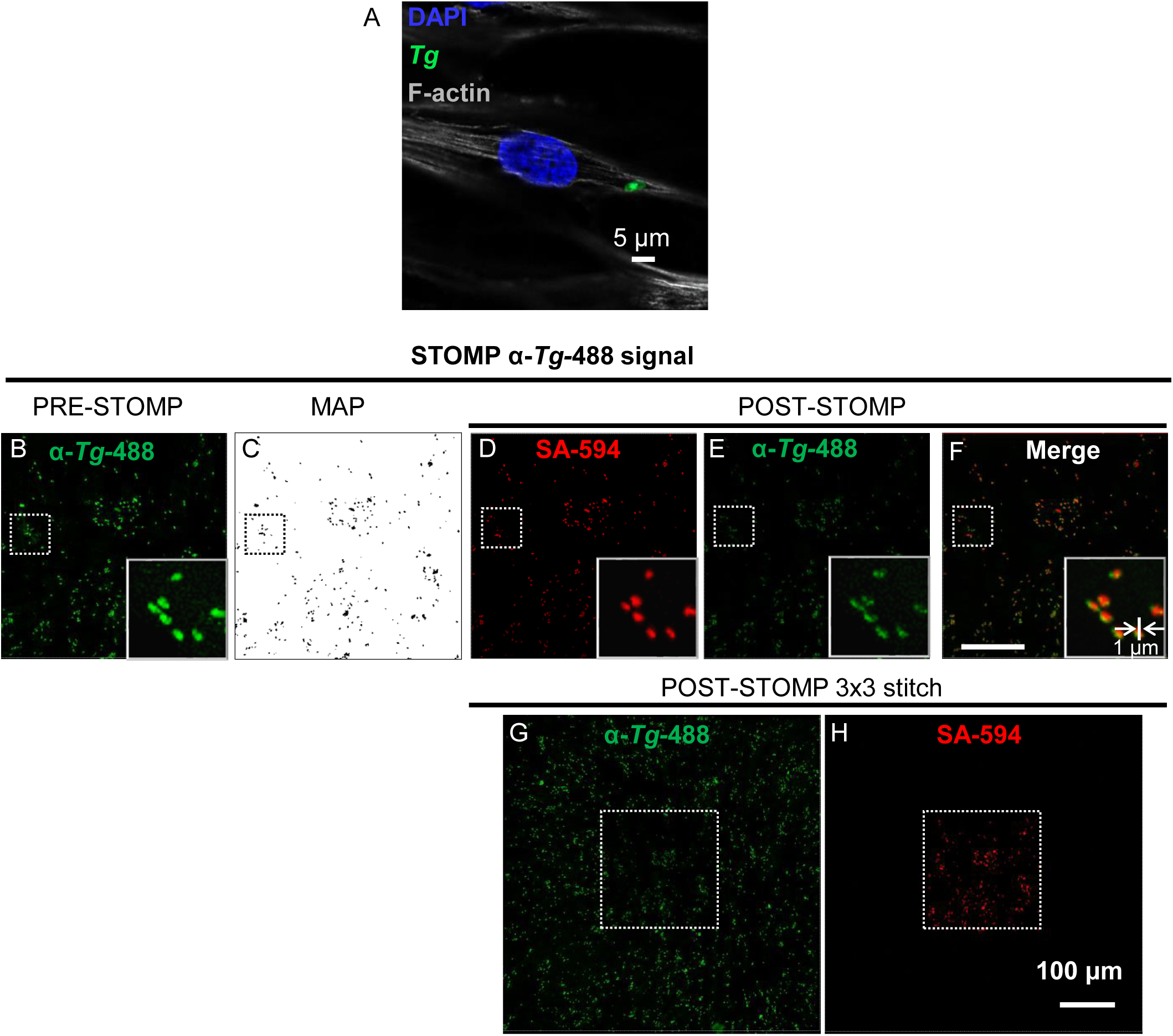
Structure of interest (SOI) proteins are selectively biotinylated by autoSTOMP UV-cross linking. **A**, Representative image of a GFP-expressing strain of *Toxoplasma* (*Tg*) infecting a human foreskin fibroblast (HFF), nuclei stained with DAPI, F-actin stained with phalloidin-660 after PFA fixation. **B**, Methanol fixed *Tg* infected HFFs were stained with a *Tg-*specific antibody directly conjugated to Alexafluor-488 (α-Tg-488, green) to identify *Tg* as the SOI. **C**, The MAP file generated in FIJI identifying the *Toxoplasma* SOI imaged in **B. D-F**, After UV-mediated biotin-BP tagging, biotinylated proteins were visualized using streptavidin-Alexafluor594 staining (**D**, SA-594, red), and directly compared to the *Tg* signal (**E**, α-Tg-488, green). Note photobleaching due to UV-targeting. **F**, merge of **D** and **E. G-H**, 3×3 tile array centered on the field of view targeted by the STOMP macro (white dotted line box): *Tg* staining (**G**, α-Tg-488, green) and biotinylation (**H**, SA-594, red). Representative of 3 independent experiments.

### AutoSTOMP biotinylation and precipitation procedures enrich for SOI proteins

To validate the efficiency of SOI biotinylation and streptavidin purification, *Tg* infected HFFs were stained and biotinylated as described in Figure 2 (Figure 3, α-*Tg* STOMP). In parallel, a negative control sample was prepared and mounted in biotin-BP, but not exposed to UV-light (Figure 3, Dark). After photo-crosslinking, samples were washed of excess biotin, dissociated from the slide in urea lysis buffer. Half of each sample was reserved as an “INPUT” control; in the other half, biotinylated proteins were enriched on streptavidin magnetic beads (Figure 3A). Both α-*Tg* STOMP and Dark samples had similar amounts of *Tg* proteins in the input control (note that polyclonal antibodies raised against *Tg* STAG recognize multiple parasite proteins) (Figure 3B, “INPUT” columns). *Tg* proteins were only enriched in α-*Tg* STOMP SA-P samples not the Dark SA-P sample (Figure 3B). In comparison, human GAPDH was detected in both INPUT samples but was not enriched in either the α-*Tg* STOMP SA-P samples or the Dark SA-P sample (Figure 3C). Probing the blots with Streptavidin conjugated to HRP confirmed that biotinylated proteins were present in the α-*Tg* STOMP samples not the Dark samples (Figure 3D). Moreover, streptavidin precipitation recovered the majority of biotin signal in the α-*Tg* STOMP SA-P sample relative to INPUT (Figure 3D). These results confirm that autoSTOMP effectively conjugates biotin-BP to SOI proteins and that biotinylated SOI proteins are enriched by the SA-P procedure.

**Figure 3.**
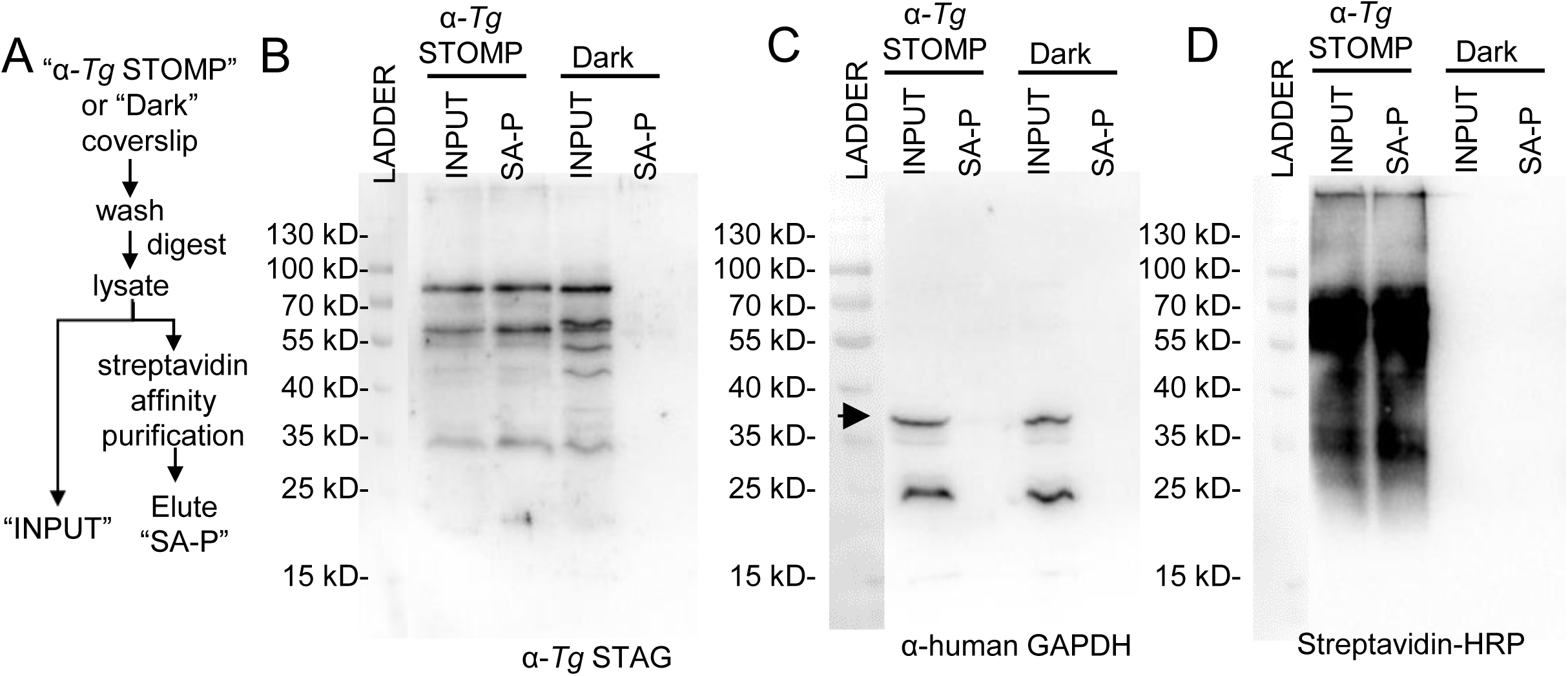
Structures of interest (SOI) proteins are enriched by autoSTOMP mediated-biotinylation and streptavidin (SA) precipitation. **A**, Schematic of the precipitation procedure. **A-D**, *Toxoplasma gondii (Tg)* infected human foreskin fibroblasts (HFFs) were stained with an α*-Tg488* antibody to identify SOI for auto-STOMP (α*-Tg* STOMP) as described in Figure 2. Alternatively, samples that have been treated identically but not exposed to UV light (Dark) serve as a control for background biotinylation and/or non-specific binding to SA-beads. Following UV crosslinking coverslips are washed to remove excess unconjugated biotin-BP, digested in 8 M urea lysis buffer and split in half. One half is reserved as the input loading control (INPUT). In the other half, biotinyated proteins are enriched by streptavidin (SA) bead precipitation. Following precipitation, samples are washed extensively and eluted in SDS Page loading dye by boiling “SA-P”. **B**, *Tg* proteins are enriched in the α-*Tg* STOMP sample relative to the Dark control when α-*Tg*488 is used to identify SOI. Note, antibody detects multiple *Tg* proteins. **C**, Human GAPDH (arrow heads) is not enriched by STOMP when α-*Tg*488 is used as a SOI. **D.** Biotinylated proteins are enriched in STOMP samples relative to input and Dark controls. Representative of 3 independent experiments.

### AutoSTOMP enriches for SOI proteins by LC-MS

We next asked if *Tg* SOI proteins could be enriched and identified by LC-MS using autoSTOMP. Samples were prepared and precipitated as described in Figure 3. After precipitation, peptides were digested on SA-beads with trypsin/LysC for mass spectrometery (Figure 4). For each sample, approximately 1 μg of protein was recovered for LC-MS. Peptide mass was identified by ion trap mass spectrometry coupled with liquid chromatography and searched against the *Tg* protein database (combined from ME49, GT1, VEG, and RH parasite strains) or the human protein database. Peptides that could not be uniquely assigned to either proteome database were excluded from downstream analysis of protein abundance. Proteins were quantified by the number of spectral counts and ranked in order of abundance (Table S-1, Figure 4).

**Figure 4.**
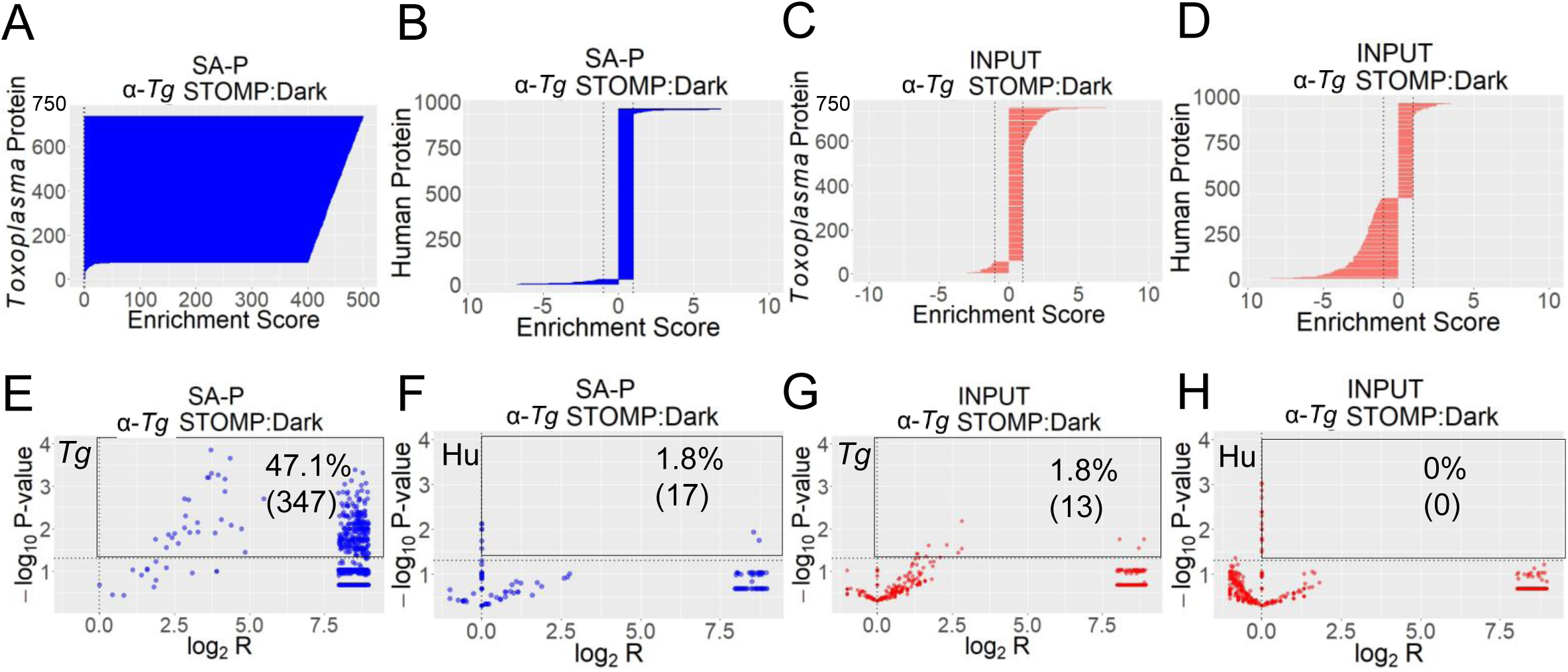
Structure of interest (SOI) proteins are enriched by peptide sequencing using auto-STOMP. *Toxoplasma gondii (Tg)* infected human foreskin fibroblast (HFF) were biotinylated and precipitated as described in Figure 3 including *Tg* SOI biotinylated by autoSTOMP (α-*Tg* STOMP) or identical samples not exposed to UV-light (Dark control). Streptavidin precipitate (SA-P) samples were on-bead digested with trypsin/lysC and the input controls (INPUT) were trypsin digested. 736 *Tg* proteins and 960 human proteins were detected with FDR < 1% for peptide and protein identification. **A-D**, *Tg* proteins were enriched in α-*Tg* STOMP SA-P samples relative to the Dark SA-P controls (**A**) whereas human proteins were not enriched (**B**). Neither *Tg* proteins (**C**) or human proteins (**D**) were enriched in the α-*Tg* STOMP INPUT samples relative to dark INPUT controls. A positive enrichment score represents the spectral count ratio (SCR) of α-*Tg* STOMP over Dark; a negative enrichment score represents the negative inverse SCR (−1/SCR) of α-*Tg* STOMP over Dark. Lines represent enrichment score of −1 and 1. **E-H**, Volcano plots representing the significance (-log_10_ P value) versus fold enrichment (log_2_ of SCR) in α-*Tg* STOMP samples relative to Dark control samples. A majority of *Tg* proteins identified are significantly enriched in α-*Tg* STOMP SA-P samples relative to Dark SA-IP (**E**, black square, significant enrichment with P < 0.05, n=3) whereas human proteins are not significantly enriched (**F**). Neither *Tg* proteins (**G**) or human proteins (**H**) are significantly enriched in α-*Tg* INPUT samples relative to Dark INPUT samples. Each dot presents a protein. Dotted lines represent y = −log_10_0.05 (significant enrichment, P < 0.05, n = 3) and x = 0 (no fold change). P-value represents student’s t test comparing each pair of samples across three independent experiments. All proteins are listed in Table S-1.

In total, 736 parasite proteins and 960 human proteins were detected in all samples with false discovery ratio (FDR) < 1% for peptide identification and an FDR < 1% for protein identification (Table S-1). The majority of *Tg* proteins identified were enriched in the α-*Tg* STOMP precipitate (SA-P) sample relative to Dark SA-P samples by spectral count ratio (Figure 4A). As expected, human proteins were not similarly enriched in α-*Tg* STOMP SA-P relative to dark SA-P (Figure 4B). In the INPUT controls, the spectral counts ratio for most *Tg* proteins (Figure 4C) and human proteins (Figure 4D) were close to 1, indicating consistent representation of proteins across samples. To determine if any of the proteins identified were significantly enriched in α-*Tg* STOMP relative to Dark samples we calculated the −log_10_ p-value over the log scale of ratio of spectral counts of the samples (Figure 4E-H). 47.1% of the 736 *Tg* proteins were significantly enriched (p-value < 0.05, a −log_10_ p-value > 1.30 and log_2_R >1) in α-*Tg* STOMP SA-P relative to the Dark SA-P sample (Figure 4E, black box). By contrast, only 1.8% of the 960 human proteins were significantly enriched in α-*Tg* STOMP SA-P relative to the Dark SA-P samples (Figure 4F). As expected, few *Tg* proteins (Figure 4G, 1.8%) and no human proteins (Figure 4H) were significantly enriched in α-*Tg* STOMP INPUT relative to and Dark INPUT controls.

We also examined protein enrichment based on abundance as the primary criterion. We considered proteins with spectral counts ≥ 5 to be represented within a high confidence interval based on the spectral count distribution model.^14^ Of the 69 high confidence *Tg* proteins in the input control 61 (88.4%) were significantly enriched in the SA-P sample (Table S-2, sheet 1; Figure S-3A-B), indicating that STOMP reliably reports the most abundant parasite proteins in the sample. Of the 667 low abundance *Tg* proteins in the INPUT controls (spectral counts < 5) 149 were represented with high confidence in α-*Tg* STOMP SA-P samples (spectral counts ≥ 5) (Figure S-3B). The 42.88% of these (Figure S-3C) were significantly enriched in the α-*Tg* STOMP SA-P sample relative to Dark SA-P sample confirming that autoSTOMP also enriched low confidence *Tg* proteins (Table S-2, sheet 2). The limited number of proteins with spectral counts ≥ 5 is likely a function of the small sample concentration used for LC-MS analysis and the sensitivity of the Thermo Scientific LTQ used in this experiment. In summary, theses data indicate that SOI proteins represented in a total cellular proteome at both high and low confidence can be tagged and detected by autoSTOMP with high confidence.

### AutoSTOMP selectively enriches proteins structures adjacent to the stained targets

An advantage of autoSTOMP compared to proximity-biotinylation methods is that the SOI can be determined by simply modifying the MAP file.^4,15^ The region surrounding the parasite is a biologically relevant organelle called the parasite vacuole membrane (PVM) that is composed of host-derived lipids, secreted parasite effector proteins, as well as host proteins that regulate vesicular trafficking, immune functions and organelle recruitment (including endoplasmic reticulum, mitochondria and lysosomes).^16^ The original STOMP protocol was shown to biotinylate structures measuring 1 μm by microscopy, however, the majority of β-amyloid plaques examined measured between 10-50 μm in diameter.^1^ We reasoned that generating a MAP file from the 1 μm region surrounding but excluding the parasite signal would allow us to identify candidate host and parasite proteins belonging to the PVM. To do this, mouse bone marrow derived dendritic cells (mBMDCs) were infected with *Tg* for two hours and samples were prepared for autoSTOMP (Figure 5A). We first confirmed that *Tg* within mBMDCs were selectively and efficiently targeted by autoSTOMP using the parameters defined in Figure 2 (Figure S-4, α-*Tg*-488). To target the region surrounding but excluding *Tg* the SOI was defined by identifying the perimeter of the α-*Tg*-488 signal (Figure 5B) and dilating the perimeter to generate a 1 μm “DONUT” outside the parasite using the “donut macro” in FIJI (Figure 5C). AutoSTOMP selectively biotinylated the DONUT SOI surrounding *Tg* at approximately 1 μm in the x and y axis (Figure 5D-F) within the targeted field of view (Figure 5G-H).

**Figure 5.**
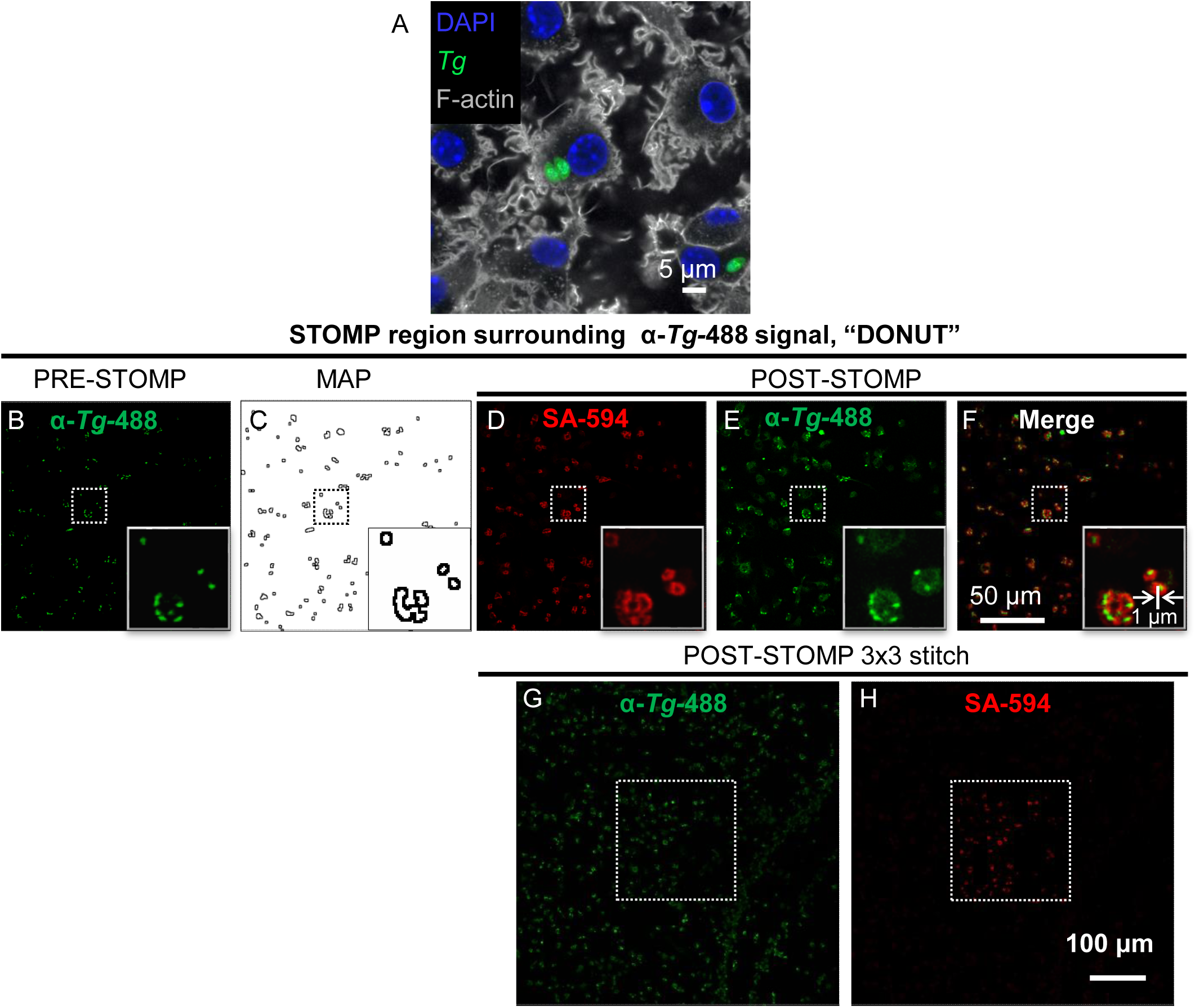
AutoSTOMP selectively biotinylates custom structure of interest (SOI) proteins. **A**, Representative image of a GFP-expressing strain of *Toxoplasma* (*Tg)* in infected mouse bone marrow derived dendritic cells (mBMDCs). Nuclei stained with DAPI and F-actin stained with phalloidin-660 after PFA fixation. **B**, Methanol fixed *Tg* infected mBMDCs were stained with a *Tg-*specific antibody directly conjugated to alexafluor-488 (α-*Tg*-488, Green) to identify*Tg*. **C**, The MAP file generated in FIJI identifying the 2 pixels surrounding but excluding the *Tg* as the SOI (DONUT). **D-F**, After UV-mediated biotin-BP tagging, biotinylated proteins were visualized using streptavidin-Alexafluor594 staining (**D** SA-594, red), and directly compared to the *Tg* signal (**E**, α-*Tg-*488, green); **F**, merge of **D** & **E**. Inset is an enlarged view of the dotted line box. **G-H**, 3×3 tile array around the STOMP targeted field of view (white dotted line box):*Tg* staining (**G**, α-*Tg-*488, green) or biotinylation (**H**, SA-594, red). Representative of 3 independent experiments.

Next, the DONUT SOI or Dark controls were precipitated as described in Figure 3 and submitted for mass spectrometry using a high-performance Velos Orbitrap LC-MS for downstream analysis (Figure 6). 138 *Tg* proteins were identified in the STOMP DONUT SA-P sample (Figure 6A), 47.1% (65 proteins) of these were significantly enriched (Figure 6B). Among the enriched *Tg* proteins were several secreted parasite proteins with known localization to the PVM or the adjacent intravacuolar network. These included rhoptry neck protein RON2, rhoptry protein ROP7, dense granule protein GRA3, microneme proteins MIC4 (Figure 6C).^17–19^ Although differences in the MS protocol mean that we cannot directly compare α-*Tg* STOMP SA-P (Figure 4E) with STOMP DONUT SA-P (Figure 6B) it is notable that the majority of parasite proteins identified within the *Tg* SOI (Table S-2) are not represented in the DONUT SOI (Table S-4), consistent with the expected spatial selection of parasite proteins trafficked to the PVM.

**Figure 6.**
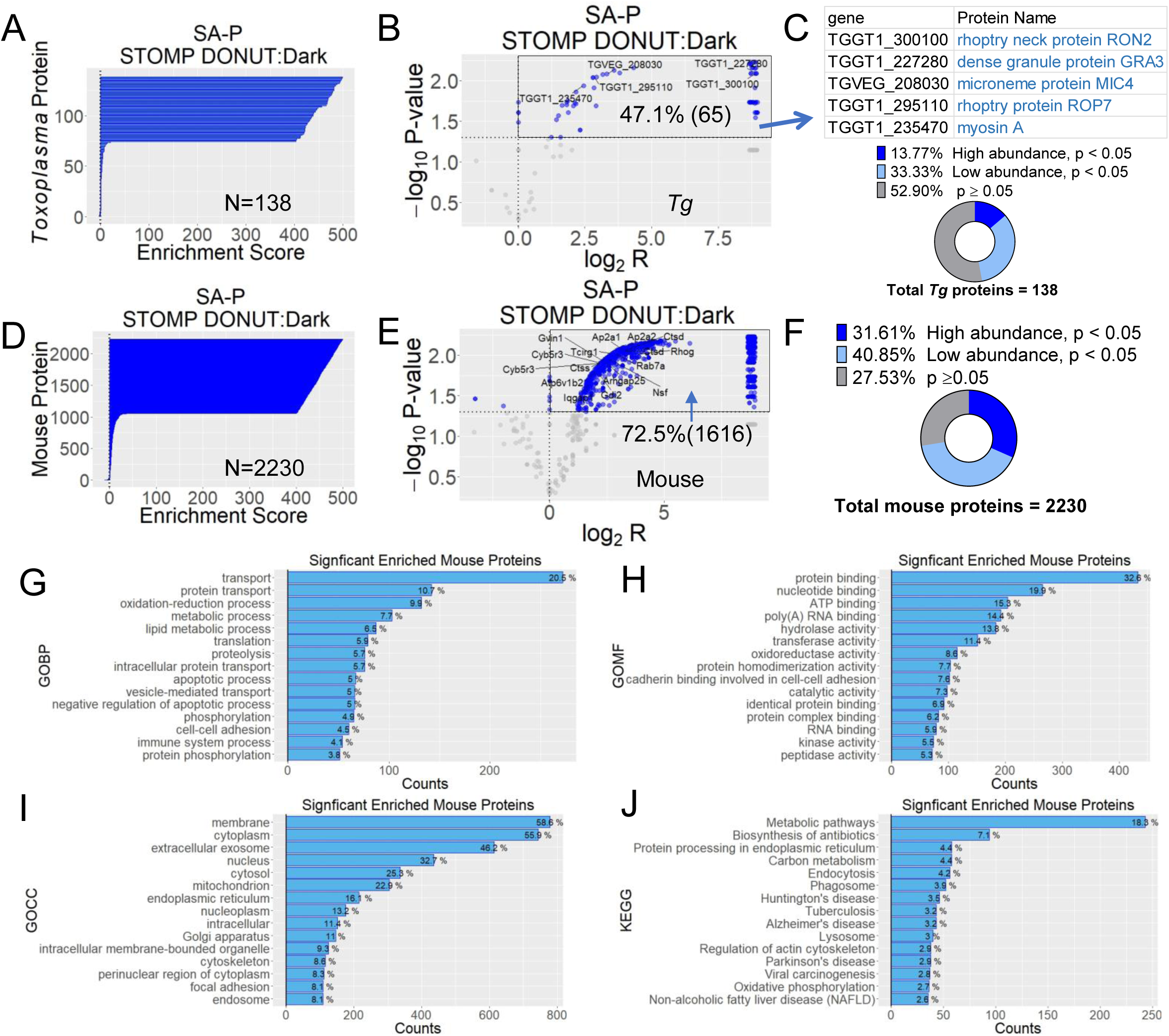
AutoSTOMP selectively identifies host and parasite proteins in custom SOI surrounding *Tg*. The regions surrounding but excluding *Toxoplasma gondii (Tg)* (STOMP DONUT) in mouse bone marrow derived dendritic cells (mBMDCs) were identified as SOI using the autoSTOMP ‘donut macro’ and biotinylated as described in Figure 5. STOMP DONUT and identical samples not exposed to UV-light (Dark) were purified by streptavidin precipitated (SA-P) and on-bead digested for LC-MS. 138 *Tg* proteins and 2230 mouse proteins were detected with FDR < 0.2% of peptide identification and FDR < 2% of protein identification. P-values are calculated from the Student’s t-test between the three replicates of STOMP DONUT SA-P sample and Dark SA-P sample (significant difference, p < 0.05; no significant difference, p ≥ 0.05). The proteins in STOMP DONUT SA-P sample with spectral counts ≥ 5 are annotated as “high abundance” whereas those with spectral counts < 5 are annotated as “low abundance”. **A-C**, Of the 138 *Tg* proteins detected in the STOMP DONUT SA-P samples relative to the Dark SA-P samples **(A)**, 47.1% or 65 *Tg* proteins were significantly enriched (**B**, blue dots). Confirmed PVM-localized proteins are indicated by their gene code and annotated in the table bellow. **C**, The 138 *Tg* proteins grouped by p-value and abundance. **D-F**, Of the 2230 mouse proteins enriched in the STOMP DONUT SA-P samples relative to the Dark SA-P samples **(D)**, 72.5% or 1616 were significantly enriched, (**E**, blue dots). A selection of host PVM associated proteins are annotated by gene name. **F**, The 2230 *Tg* proteins grouped by p-value and abundance**. A&D**, A positive enrichment score represents the spectral count ratio (SCR) of STOMP DONUT over Dark; a negative enrichment score represents the negative inverse SCR (−1/SCR) of STOMP DONUT over Dark. **B&E** Volcano plots show the significance (-log_10_ P value, n = 3) versus fold enrichment (log_2_ of SCR, mean of three replicates) in STOMP DONUT samples relative to Dark samples. Blue indicates p < 0.05, or gray indicates p ≥ 0.05. Black box indicates proteins significantly enriched by autoSTOMP. **C&F**, Proteins enriched in the STOMP DONUT sample relative to the Dark control are categorized by p-values and spectral counts above ≥ 5 (high abundance) or below < 5 (low abundance) represented as pie charts. N= 3 independent experiments. **G-J**, The 72.5% or 1616 significantly enriched mouse proteins (**E**) are plotted with the Gene Ontology (GO) terms or KEGG terms against the counts of the proteins involved. Three types of (GO) terms (Biological Process (BP, **G**), Molecular Function (MF, **H**), or Cellular Component (CC, **I**) or KEGG pathway terms (**J**) are used. The terms are ranked by the counts from the largest to the smallest and annotated with the percentage of involved proteins out of the total. Here, only the top 15 terms are shown.

2230 mouse proteins were identified in the STOMP DONUT SA-P relative to Dark SA-P (Figure 6D), 72.5% of which were significantly enriched (Figure 6E). In comparison to the α-*Tg* STOMP SA-P proteome in which significantly enriched parasite proteins were 19 times more abundant than host proteins (Figure 4E-F), the STOMP DONUT SA-P contained 24 times the number of significantly enriched host proteins compared to *Tg* proteins (Figure 6E&B). At least 16 of these host proteins have been detected at the PVM or the intravacuolar network during *Tg* infection (Figure 6E). Ras-related protein Rab-7 and cathepsin D are associated with autophagosomal formation around PVM and lysosomal fusion *Tg* clearance.^16,20^ The AP-2 regulates endosomal vesicle fusion with autophagosomes and the PVM.^21^ Proteins expressed in the mitochondria including the V-type proton ATPase and Rab associated proteins that regulate Golgi vesicular transport might also be expected based on reports of recruitment of these structures to the PVM.^21–23^

In addition to identifying host proteins that have been experimentally confirmed at the PVM, we searched the significantly enriched mouse proteins against gene ontology (GO) and Kyoto Encyclopedia of Genes and Genomes (KEGG) pathway annotation libraries with David Bioinformatics Resources 6.8.^12^ Using the GO term “biological processes”, 20.5% of STOMP DONUT SA-P proteins belong to the “transport” annotation group and 7.7% are in “metabolic processes” annotation group (Figure 6G) consistent with parasite scavenging from the host.^24^ Using the GO term “molecular function,” 32.6% proteins are associated with “protein binding” and 15.3% are involved in ATP binding consistent with regulation of membrane fusion at the PVM (Figure 6H).^25^ Using the GO term cellular component, more than 55% proteins are annotated with “membrane” GOCC term (Figure 6I). KEGG analysis also indicated that 18.3% of proteins belong to the “metabolic pathways” annotation (Figure 6J); and proteins associated with organelles that are known to be recruited to the PVM were represented including endolysosomal vesicles, endoplasmic reticulum and mitochondria.^16,25–27^ Based on these data, we conclude that autoSTOMP selectively enriches proteins belonging to sub-cellular regions with spatial resolution of approximately 1 μm.

## CONCLUSION

Here we show that autoSTOMP is a fully automated platform to visualize, tag, affinity purify and identify proteins by mass spectrometry. AutoSTOMP represents a significant technological advance because it liberates the user from field-by-field task management previously required to perform STOMP. In automating the STOMP protocol, we retained and refined the ability to define regions of interest based on structure size, shape or localization using FIJI image analysis software. This flexibility is a major advantage of STOMP compared to other proximity-based tagging tools ^2,4^

Our choice of a biotin-BP was based on the low background reactivity of the BP group and the high affinity of the biotin-streptavidin interaction. Benzophenone covalently inserts into N-H or C-H bond upon excitation with UV-wavelength light^1^. However, the cost of this tagging strategy is that most BP modified peptides will remain attached to the SA beads. Any eluted peptides bearing biotin-BP will not be recognized by MS, due to the uncertain mass shift imparted by the tag. For this reason, an ideal ratio of biotin-BP would be one tag per protein which may be optimizable in future experiments.

Finally, it is important to note that autoSTOMP is not limited to host-pathogen interactions as studied here. We chose this system because it took advantage of organisms with diverse and divergent proteomes in extremely close proximity to one another, which allowed us to accurately quantify protein enrichment in SOIs relative to background. Our data suggest that autoSTOMP can be used to identify proteins belonging to 1 μm scale sub-cellular structures in a range of biological specimens.

## ASSOCIATED CONTENT

### Supporting Information

1. Table S-1. The *Tg* and Human proteins identified in the samples from *Tg* infected HFFs related to Figure 4.
2. Table S-2. The *Tg* proteins that are filtered under two different conditions related to Figure 4: “INPUT > 4”: proteins beyond the SC cutoff of 5 in the INPUT controls; “SA-P > 4 INPUT < 4”: proteins below the SC cutoff of 5 in the INPUT controls, but beyond the SC cutoff of 5 in the SA-P samples.
3. Table S-3. The *Tg* and mouse proteins identified in the *Tg* surrounding regions excluding *Tg* in mBMDCs infected with *Tg* related to Figure 6.
4. Table S-4. The *Tg* and mouse proteins significantly enriched in the *Tg* surrounding regions excluding *Tg* related to Figure 6.
5. Figure S-1. The photo-crosslinking condition was optimized at power/iteration/dwell time in ms (P/I/D) = 3/1/1.
6. Figure S-2. Optimize the biotin-BP at a concentration of 1 mM for photoactive labelling.
7. Figure S-3. Half of total *Tg* proteins with either high or low abundance (confidence) in the INPUT controls were enriched in α-*Tg* STOMP SA-P samples by autoSTOMP procedure (related to Figure 4E).
8. Figure S-4. Validate the photo-crosslinking biotinylation on *Tg* SOIs in mBMDCs infected with *Tg* with autoSTOMP.

## Author Contributions

B.Y., R.R. and S.E.E. designed the experiments in this study. X.Z. provided biological specimens for these studies. B.Y. performed the STOMP experiments, data processing and analysis. R.M. and K-L.H conducted Mass Spec experiments using the Thermo Scientific LTQ. B.Y. and S.E.E. prepared the manuscript.

## Notes

The authors declare no competing financial interest.

## ACKNOWLEDGMENTS

We thank Dr. Nicholas E. Sherman and Dr. Jeong-Jin Park at the W.M. Keck Biomedical Mass Spectrometry Laboratory at the University of Virginia’s School of Medicine for providing Mass Spectrometry analysis service for experiments using the Velos Orbitrap. We thank Anson Parker at the University of Virginia School of Medicine Library for the introduction to SikuliX. We thank Dr. Hardik at the University of Virginia School of Medicine, for assistance in the design and execution of the Python script to process *Tg a*nd human protein assignments. We thank Dr. Josh Elias at The Chan Zuckerberg Institute for guidance in the design and execution of mass spectrometry experiments. We are grateful to Dr. Avi Chakraborty and Dr. Kevin Hadley at the University of Toronto for extensive discussion of STOMP optimization, initial training with the STOMP pipeline and assistance using the STOMP Macro.

This work was supported by NIH K22 AI116727 (SEE) AMH/Allen Institute 31315 (SEE) and start-up funds from the University of Virginia SOM and the Emily Couric Cancer Center.

---

**Figure S-1.**
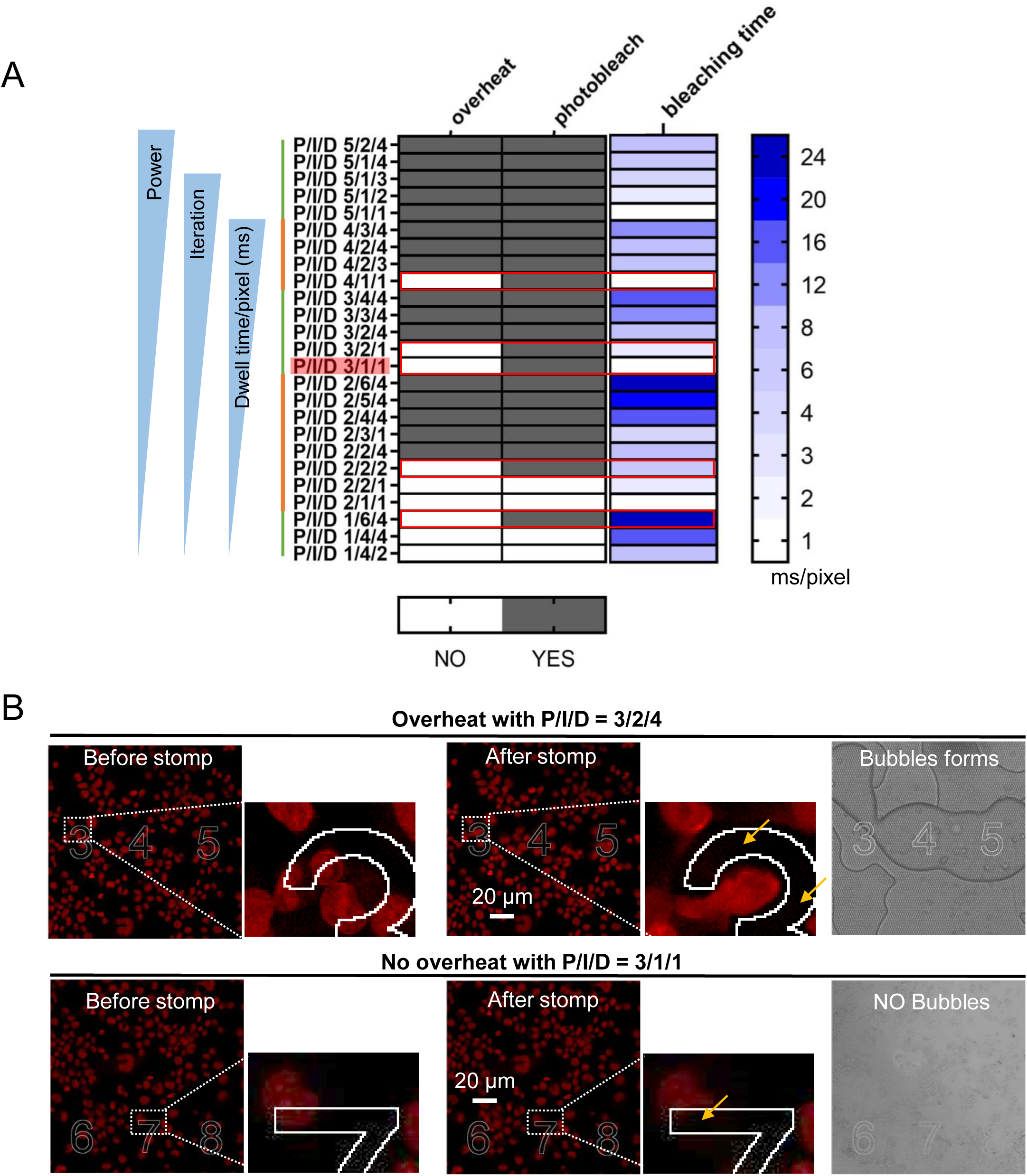
The optimal photo-crosslinking condition was determined varying the laser power (P), the number of iterations each field of view was photo-excited (I) and the dwell time (D) represented as “P/I/D” using methanol fixed THP-1 cells stained with propidium iodide. **A**, Five conditions were identified with sufficient power for photo-bleaching that did not induce overheating (red box). P/I/D = 3/1/1 was chosen to minimized the time necessary to process each field of view. Samples are primarily ordered by the power (P), secondly ordered by the iteration time (I) and thirdly ordered by the Dwell time/pixel (ms). The bleaching time per pixel in milliseconds (ms) to excite the photo-crosslinking equals the product of the iteration time and Dwell time/pixel. **B**, Representative laser settings for photo-crosslinking that causes the overheating with P/I/D = 3/2/4 (Top) or no overheating with P/I/D = 3/1/1 (Bottom) inside the slide. The STOMP crosslinking was directed to the SOIs with the shapes of numbers “3”, “4”, and “5” (Top) or numbers “6”,”7”,”8” (Bottom). The photobleaching occurs in the auto-STOMP targeted SOIs (yellow arrow) while the outside area was not affected (outside the white outline). Bubbles formed were detected under the bright field indicating overheating.

**Figure S-2.**
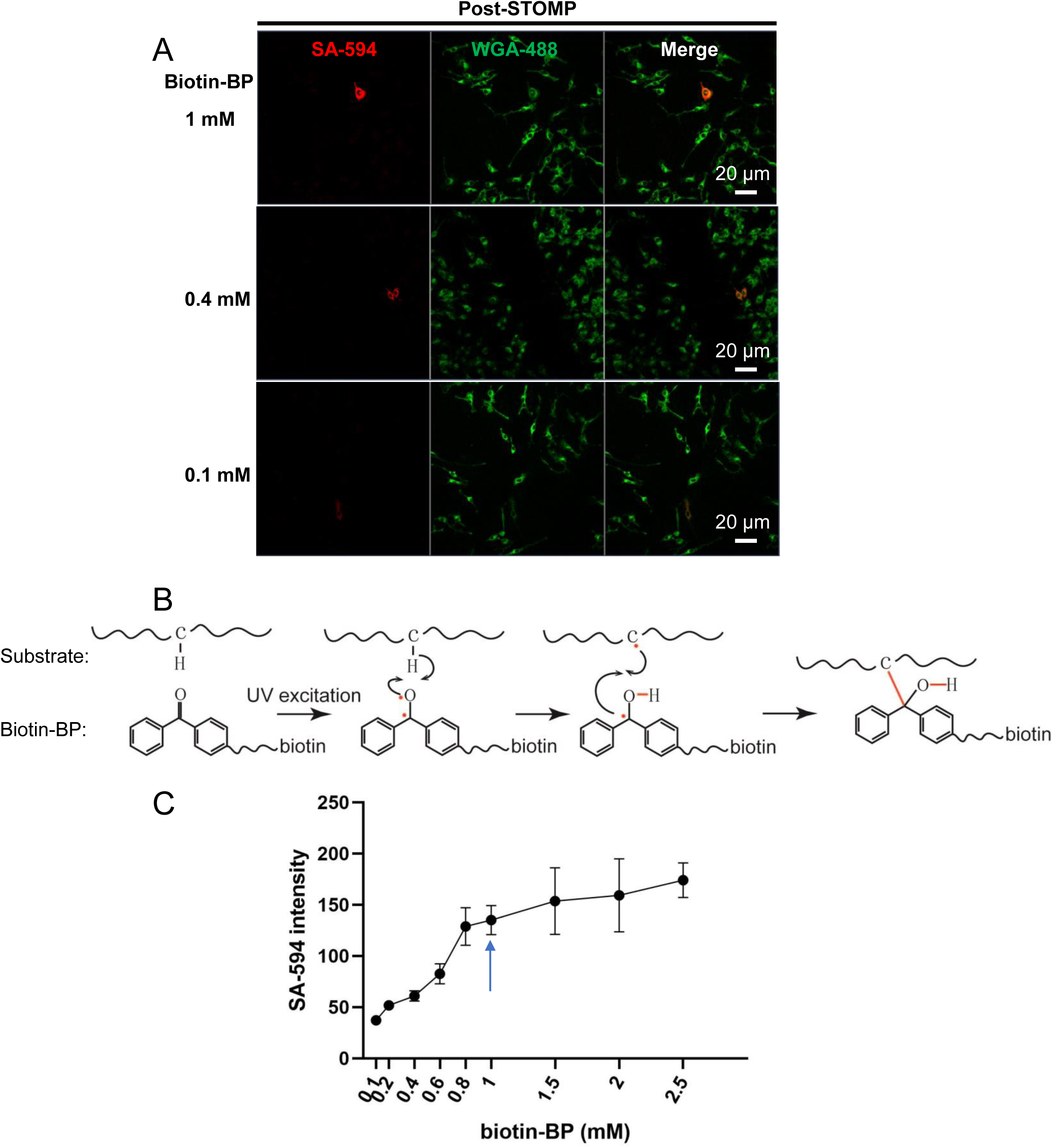
Biotin-BP concentration was optimized by titration 0.1mM to 2.5mM biotin-BP in mounting media. Immortalized mouse bone marrow derived macrophages (iBMDMs) were methanol fixed and stained with fluorescein labeled wheat germ agglutinin (WGA-488) then mounted with biotin-BP at concentrations of 0.1 – 2.5 mM. Samples were photo-crosslinked at 720 nm at laser Power/Iteration/Dwell time = 3/1/1. **A**, Representative images of iBMDMs (green) and 0.1, 0.4 and 1mM biotin-BP showing an individual cell targeted with streptavidin-594 (red). **B**, A diagram shows biotin-BP inserts into C-H or N-H bonds under UV. **C**, SA-HRP staining relative to biotin-BP concentration. 1mM biotin-BP (blue arrow) was selected as the lowest concentration needed for optimal SA-594 staining.

**Figure S-3.**
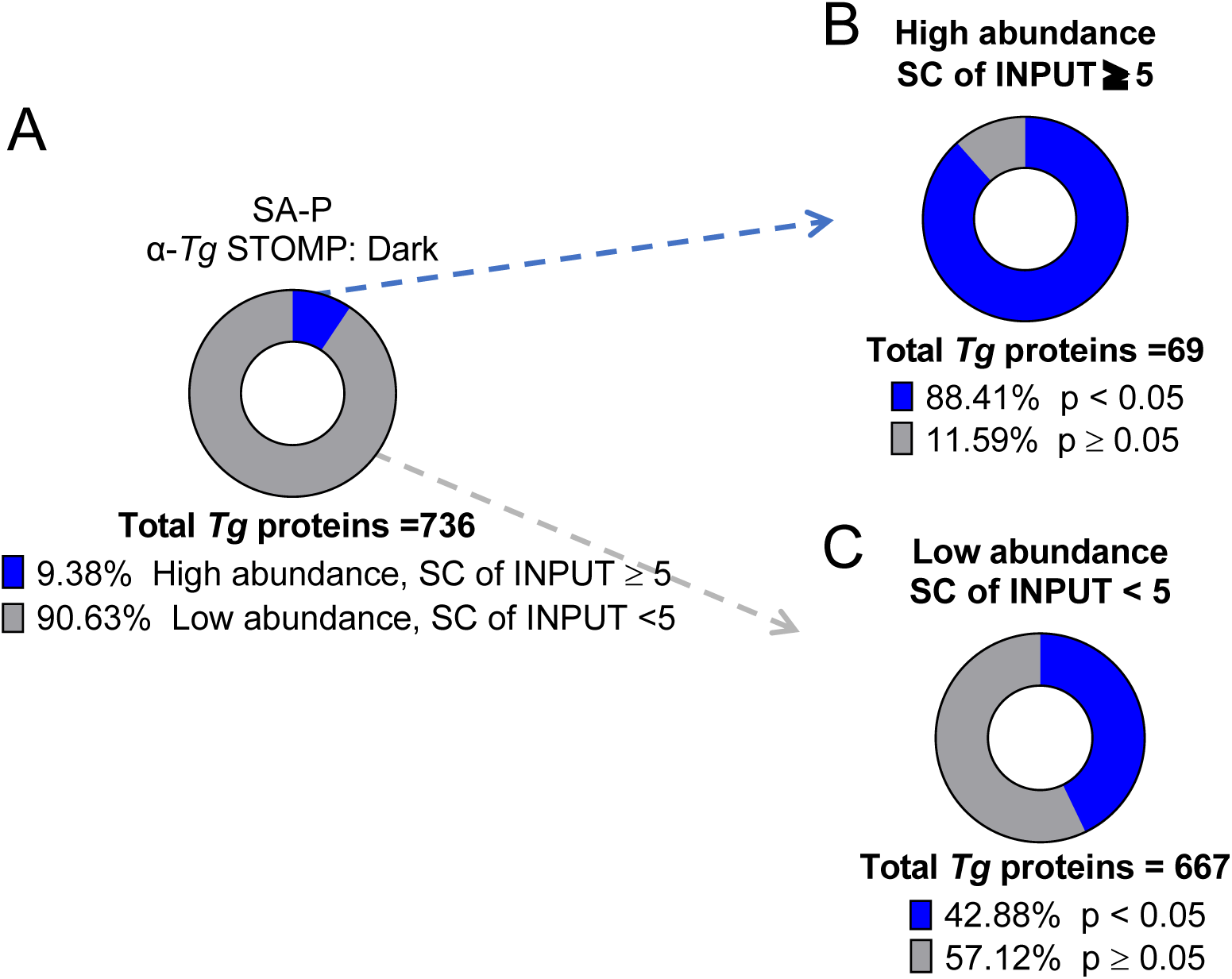
Both high and low abundance proteins are enriched by autoSTOMP. **A**, Of the 736 *Toxoplasma gondii (Tg)* proteins identified in Figure 4E, 9.38% of proteins are “high abundance” with spectral counts (SC) in the INPUT controls ≥ 5, and 90.63% of proteins were “low abundance” with SC in the INPUT controls < 5. p-values based on student’s t-test comparing α-*Tg* STOMP SA-P and Dark SA-P samples. N=3 independent experiments (significant difference, p < 0.05; no significant difference, p ≥ 0.05). **B**, 88.41% of the high abundance proteins are significantly enriched by autoSTOMP. **C**, Whereas, 42.88% of the low abundance proteins are enriched in the α-*Tg* STOMP compared to the Dark.

**Figure S-4.**
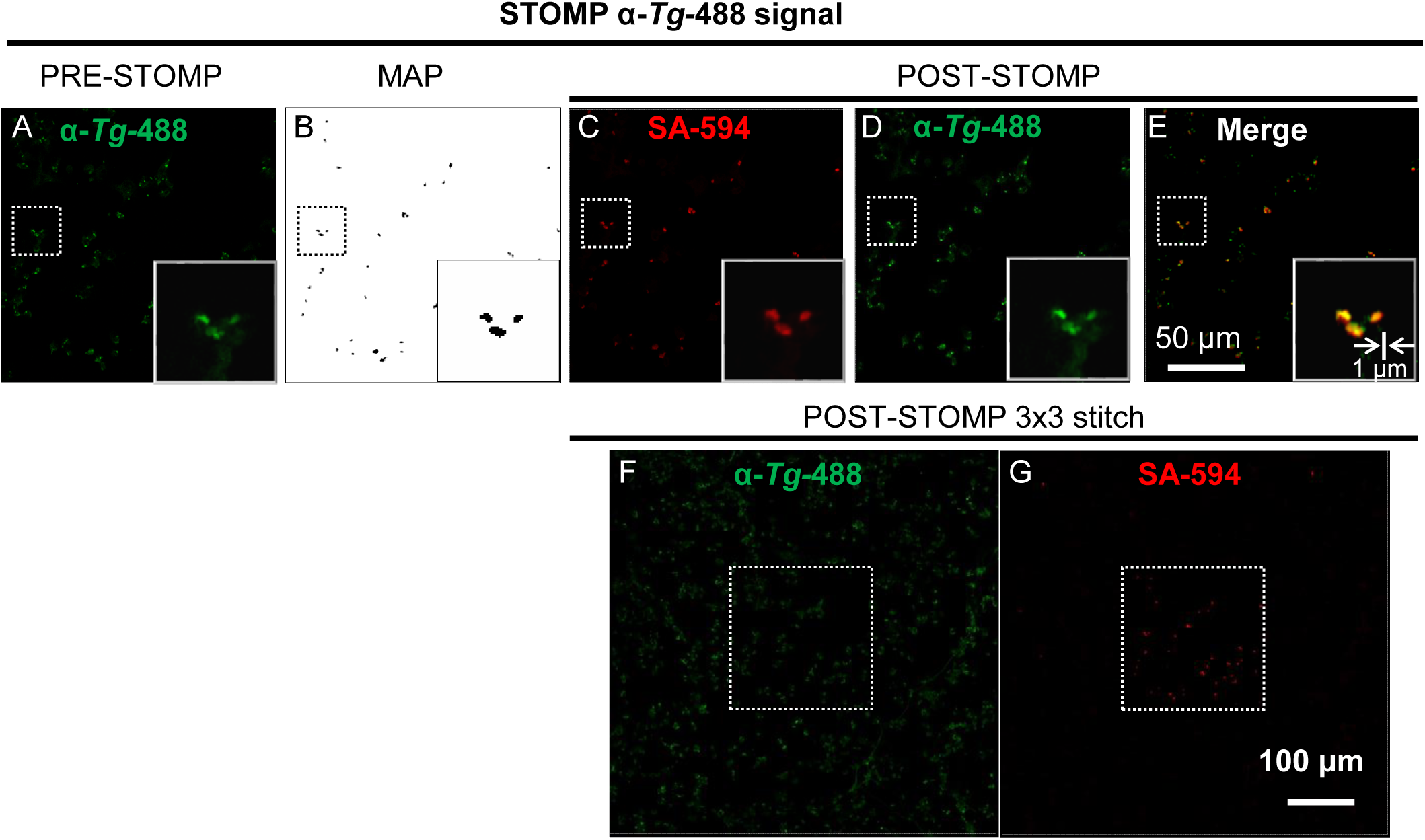
The *Tg* is selectively biotinylated by auto-STOMP UV-cross linking in *Toxoplasma* (*Tg*) infected mouse bone marrow derived dendritic cells (mBMDCs). **A**, *Tg* in mBMDCs were stained with a *Toxoplasma-*specific antibody directly conjugated to alexafluor-488 (α-*Tg-*488, green) to identify *Tg* as the SOI. **B**, The MAP file generated in FIJI identifying the *Tg* SOI imaged in **A. C-E**, After UV-mediated biotin-BP tagging, biotinylated proteins were visualized using streptavidin-Alexafluor594 staining (**C**, SA-594, red), and directly compared to the *Tg* signal (**D**, α-*Tg-*488, green). **E**, merge of **C** and **D. F-G**, 3×3 tile array centered on the field of view targeted by the STOMP macro (white dotted line box):*Toxoplasma* staining (**F**, α-*Tg-*488, green) and biotinylation (**G**, SA-594, red). Representative of 3 independent experiments.

